# Multiscale Metapopulations

**DOI:** 10.1101/2025.08.13.670192

**Authors:** Daniel Malagon, Benjamin T. Camper, Sharon Bewick

**Affiliations:** Department of Biological Sciences, Clemson University, Clemson, SC, USA

## Abstract

New advents in technology have improved our ability to study and track host-associated organisms. Often, these organisms experience vastly different spatial and temporal scales than their hosts. However, host-associated organisms are not independent of their hosts nor are hosts independent of their host-associated organisms. Rather, processes at the host and host-associated organism scales interact to alter outcomes for both the host and its symbionts. Often, host-associated organisms are described in terms of metapopulations, wherein individual hosts act as habitat patches for their symbionts. When hosts themselves also exist in patchy environments, the result is metapopulation structure at two distinct scales. In this paper, we develop a model to explore how the incorporation of multiscale metapopulation dynamics impacts metapopulation predictions at both the host and symbiont scales. While related to several disease models, our system differs in its inclusion of full colonization and extinction dynamics at both scales, and its emphasis on how the interaction between colonization and extinction at these two scales impact the host and its host-associated organisms, including pathogens as well as symbionts that are beneficial to the host. Overall, our multiscale metapopulation model predicts rich dynamics that often deviate from the dynamics of metapopulation models built for either the host or its host-associated organisms independently.

## Introduction

Richard Levins first defined a metapopulation as “a population of populations.”^1,2^ That is, a set of populations, each occupying a discrete habitat patch, with the habitat patches themselves distributed across a larger landscape. The resulting ‘patchy’ spatial structure is important because it can lead to population dynamics that are distinct from otherwise comparable ‘well-mixed’ populations. Often, for example, the long-term survival of species in a metapopulation depends on the stochastic balance between extinction and recolonization among metapopulation patches.^1-3^ In the 50 years since its origin, metapopulation theory has become a cornerstone of population, community, and landscape ecology.^4^ It has been used to explain everything from local and regional extinction processes,^4-6^ to land-use planning for conservation,^7-9^ genetic structuring of endangered species,^10^ and even disease dynamics.^11-13^

Metapopulations are often hailed as a classic example of the incorporation of spatial scale into ecology.^14^ However, while metapopulations do include distinct local and regional scales, local populations and regional metapopulations are each considered at their own, typically arbitrary, spatial scale. Indeed, even classifying a system as a metapopulation often depends on the *a priori* selection of a single local and a single regional scale.^15^ This is problematic because the persistence and predicted dynamics of the metapopulation often depend on the subjectivity of defining what is the local and the regional scale of interest.^15,16^ These definitions, however, are rarely obvious,^17,18^ particularly in metapopulations where patchiness exists simultaneously over multiple scales.^17^ Extending the classic metapopulation framework to ‘multiscale metapopulations’ remains an open challenge.

One particularly clear example of a multiscale metapopulation is a population of host-associated organisms distributed across hosts that are, themselves, confined to a metapopulation of habitat patches (Figure 1). This type of structure can emerge from host-associated organisms that are either macro- or microscopic, that are either endo- (e.g., gut microbe) or ectoparasitic (e.g., skin microbe), and that have effects on the host ranging from pathogenic to neutral, commensal, or even mutualistic. Thus, there are a broad range of systems that can be described by this ‘two-scale’ metapopulation organization. At the smaller of the two scales, individual hosts function as patches for host-associated organisms. Host-to-host transfer of these host-associated organisms then links macroscopic hosts into a small-scale metapopulation, while host turnover or host clearance of the symbiont acts as a constant source of small-scale patch extinction. Meanwhile, the hosts themselves inhabit and disperse between their own habitat patches in a second, much larger scale metapopulation. However, in moving between their own habitat patches, the hosts bring their host-associated organisms with them. Thus, host-associated organisms experience metapopulation structure at two scales: (1) extinction and colonization of new hosts (small-scale) and (2) extinction and colonization of new host habitat patches (large-scale).

**Fig 1.**
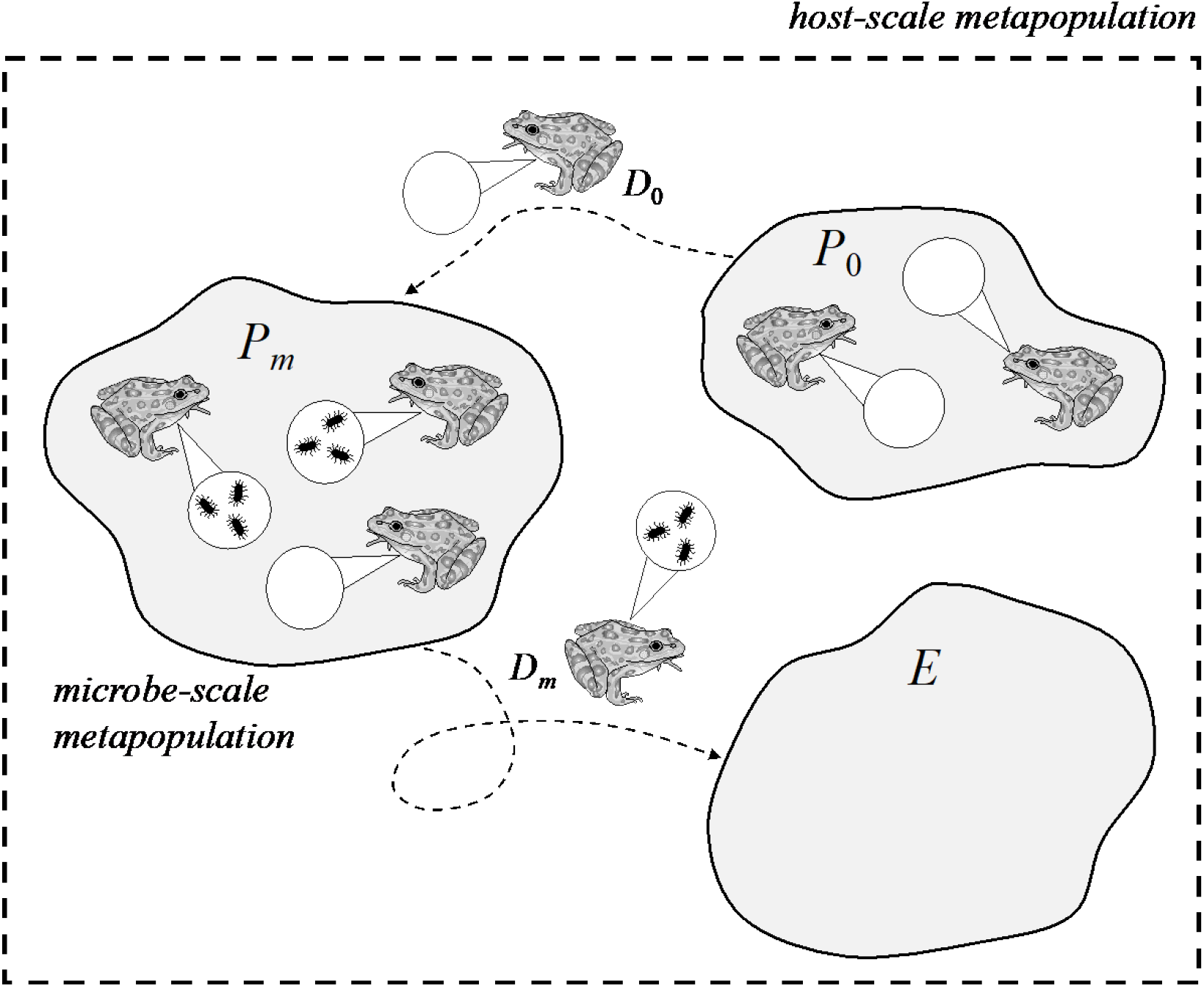
A multiscale metapopulation. Here frogs represent hosts that may or may not be colonized by a focal microbial population, while ponds represent host patches that may or may not be colonized by frogs and, by extension, may or may not be colonized by the focal microbe. *P*_*m*_, *<u>P</u>*_*<u>0</u>*_, and *E* label microbe-carrying, microbe-free, and empty patches. Hosts with (*m*) or without the microbe of interest (1*-m*). Dispersing hosts with (*D*_*m*_) and without *(D*_*0*_) focal microbe.

At this point, it is worth noting that multiscale metapopulations, and indeed metapopulations in general, share a deep connection to disease dynamics. For example, the classic Levin’s metapopulation model^19^ is mathematically equivalent to the susceptible-infected-susceptible (SIS) model – a mainstay of mathematical epidemiology. While the effects of scale remain relatively poorly explored in metapopulation literature, the issue of scale has long been a focus in disease models.^20,21^ This includes studies of ‘patchy’ SIS models^21-28^, which are, in essence, systems with partial multiscale metapopulation structure.^29^ However, while patchy SIS models capture some aspects of multiscale metapopulations, they fail to account for full metapopulation dynamics at two distinct scales. Rather, patchy SIS models consider disease spread and host recovery/death and replacement (small-scale colonization and extinction dynamics), as well as host movement between habitat patches (large-scale colonization dynamics), but generally fail to consider destruction of host habitat patches (large-scale extinction dynamics). Said differently, the focus of patchy SIS models is the effect of rare movements of susceptible and infected hosts between cities like Boston, Chicago, and New York. Rarely, however, do these models ask how disease dynamics are impacted when Boston is destroyed. Thus, in most patchy SIS models, the host does not persist as a true metapopulation. Rather, patchy SIS models describe an (micro)organism that faces small-scale metapopulation dynamics coupled to large-scale dispersal limitation of its host. Another closely related model from epidemiology that captures partial but not full multiscale metapopulation dynamics is the Hess model.^11,12^ Hess explored the dynamics of susceptible-infected (SI) models in metapopulations and, unlike most patchy SIS models, did consider catastrophic loss of host patches^11,12^. However, because Hess’ focus was on SI models, rather than SIS models, his systems incorporated both colonization and extinction at the host-scale, but only colonization at the microbe-scale. In other words, a host, once infected, remains infected indefinitely and infected patches can only become susceptible again through host patch destruction and subsequent recolonization by susceptible hosts. As a consequence, there is no requirement for the disease to overcome colonization:extinction constraints within individual host patches. Thus, Hess’ models do not, at least explicitly, capture both colonization and extinction at the microbe scale.

Beyond epidemiology, several other types of systems have been used to explore different aspects of multiscale metapopulations. The effects of multiscale dispersal, for instance, have been studied using islands comprised of multiple habitat patches.(Huth et al. 2015) In these systems, any individual habitat patch can be colonized via either within- (small scale) or between- (large scale) island dispersal, capturing colonization processes at two separate scales. Extinction, however, typically occurs at a uniform rate across individual populations (i.e., only at the smaller of the two scales); thus, like epidemiological models, these models do not consider full metapopulation dynamics at both scales. Parasite-hyperparasite interactions provide another closely related class of systems that encompass some aspects of multiscale metapopulations.(Holt and Hochberg 1998; Klinkenberg and Heesterbeek 2005; Parratt and Laine 2016; Sandhu et al. 2021) In these systems, hosts function as habitat patches, parasites as hosts and hyperparasites as host-associated organisms. Accordingly, parasite-hyperparasite systems are often modeled with sets of equations similar to the ones that we propose (see ‘Methods’). Unlike epidemiological or multi-patch island models, these systems consider full colonization and extinction dynamics at two separate scales (parasite and hyperparasite). However, the very specific nature of parasite-hyperparasite interactions means that models of parasite-hyperparasite systems have only focused on a subset of the dynamics that might emerge from multiscale metapopulations. Hyperparasites are, for instance, either neutral or pathogenic to the parasites that they infect. Consequently, parasite-hyperparasite systems have not been used to explore the impacts of beneficial host-associated organisms. Likewise, models of parasite-hyperparasite systems have not considered Allee effects among parasites. This is, nevertheless, a phenomenon that is commonly studied via the ‘rescue effect’ in metapopulation literature. How the rescue effect might modify multiscale metapopulation dynamics, even for neutral and pathogenic host-associated organisms, remains poorly understood.

In this paper, we seek to advance metapopulation theory (and, by extension, SIS models and/or SI models in metapopulations) by developing a general and fully multiscale metapopulation framework. Specifically, our goal is to extend the Levins and Hess spatially implicit metapopulation models by considering colonization/extinction dynamics that occur simultaneously across two distinct scales: the host-associated organism scale and the host-scale. In addition, we consider several other features that are typically absent from parasite-hyperparasite and epidemiology models, including the potential for the microbe to be beneficial to the host, as well as a host rescue effect. At the smaller of the two scales that we consider, we use a traditional Levin’s model. At the larger of the two scales, we use an extended system that includes 3 patch states: empty patches, patches inhabited by hosts that are not inhabitated by the host-associated organism, and patches inhabited by hosts carrying the host-associated organisms (Figure 1). This latter framework, that is akin to Hess’ approach, but that includes explicit accounting for microbe dynamics at the smaller scale, allows us to examine how the balance between colonization and extinction at two distinct scales impacts system dynamics, patch occupancy and metapopulation persistence.

## Methods

We develop a two-scale extension of the classic Levins model.^1,2,19^ For the remainder of the paper, we will refer to the smaller of the two scales as the ***microbe-scale*** and the larger of the two scales as the ***host-scale***. This is primarily for ease of reference, and there is no strict requirement that the smaller of the two scales involve a microbe. Rather, the only requirement is that small-scale dynamics involve some form of host-association. Like the Levins model, our model makes several simplifying assumptions:^19^

1. There are an infinite number of identical host patches, and all host patches are equally strongly connected to one another via dispersal/migration (i.e., mean-field approximation at the host-scale).
2. Each host patch contains an infinite number of hosts, and all hosts are equally strongly connected to one another via host-host contact/microbe transfer (i.e., mean-field approximation at the microbe-scale).
3. Host habitat patches are either occupied or unoccupied by the host and/or microbe, with no accounting for variation in the abundances of either microbe-carrying hosts or hosts overall (i.e., a presence/absence model at the host-scale).
4. Hosts either carry or do not carry the microbe, with no accounting for variation in abundance of the microbe on any given host (i.e., a presence/absence model at the microbe-scale)
5. There is a constant per patch rate of host extinction from host habitat patches; this could be due to stochastic loss of hosts from the habitat patch or complete patch destruction, followed by patch re-creation either in the same location or a new location (e.g., mowing of a field that was home to butterflies or drying of a pond that was home to amphibians).
6. There is a constant per host rate of microbe extinction from individual hosts; this may be due to active clearance of the microbe by the host immune system, stochastic loss of the microbe by the host or host turnover without vertical transmission (i.e., a microbe-carrying host dies and is replaced by a newborn microbe-free host)
7. The rate of habitat patch colonization by the host is proportional to an intrinsic constant, multiplied by both the number of occupied and unoccupied habitat patches (i.e., mass-action at the host-scale).
8. The rate of host colonization by the microbe is proportional to an intrinsic constant multiplied by the number of microbe-carrying hosts and the number of microbe-free hosts that are present in any given host habitat patch (i.e., mass-action at the microbe-scale)

In what follows, we define the basic components of our multiscale metapopulation model, along with several extensions, including a model with a rescue effect and a mainland-island model; both of which are common extensions of the classic, single-scale Levins model.

### Microbe-Scale Metapopulation Model

From the perspective of a dispersing host-associated microbe, each host is a patch capable of harboring an entire microbial population. At this scale, there are two possible states for each patch (i.e., each host) – hosts can either be microbe-carrying (occupied) or microbe-free (unoccupied). Assuming a basic Levins model gives:

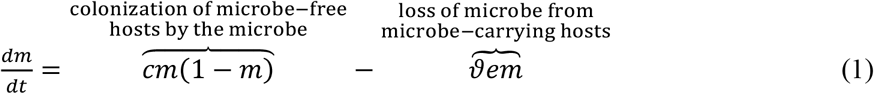

where *m* is the proportion of microbe-carrying hosts, *c* is the rate at which microbes colonizes microbe-free hosts and *e* is the rate at which microbes are lost from a host (this may be due to host clearance, stochastic loss of the microbe from the host, or host turnover without vertical transmission). Because pathogenic and mutualistic microbial associates have the potential to impact host death and, by extension, microbe-scale population extinction events, we include an additional parameter, *ϑ*, that allows the microbe to increase (*ϑ* > 1, pathogen) or decrease (*ϑ* < 1, mutualist) microbial extinction rates (via effects on host turnover). The solution to equation (1) is the well-known Levins result:

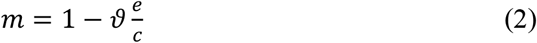

### Basic Host-Scale Metapopulation Model

At the host-scale, a patch is any region of habitat capable of supporting a breeding population of the host (e.g. pond-breeding frogs in a series of interconnected ponds, see Figure 1). Again, taking inspiration from the Levins model, we ignore host and microbe abundances within individual patches and instead treat host patches as empty (no hosts or microbes), microbe-free (hosts present, but none carry the microbe), or microbe-present (hosts present, at least some carry the microbe; see Figure 1). This allows us to define the following system of equations describing the dynamics of host patch occupancy:

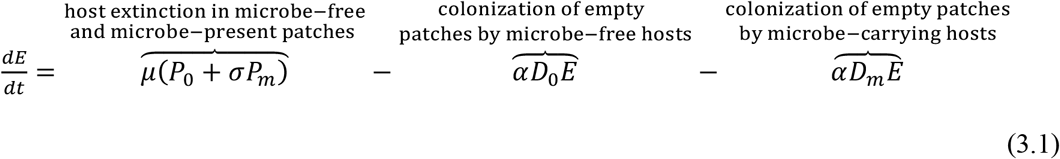

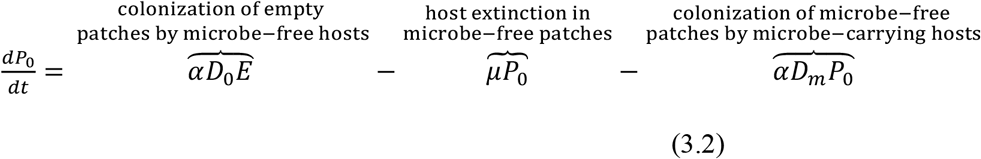

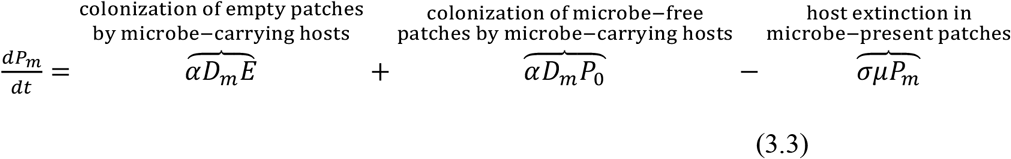

In equation (3) *E, P*_0_, and *P*_*m*_ are the proportion of habitat patches that are empty, microbe-free and microbe-present respectively, *D*_0_ and *D*_*m*_ are the number of dispersing hosts that are microbe-free and microbe-carrying respectively, *α* is the rate at which dispersing hosts successfully colonize habitat patches, and *µ* is the rate at which host extinction occurs in host habitat patches. Finally, we include the parameter *σ*, to allow for the possibility that the microbe increases (*σ* > 1, pathogen) or decreases (*σ* < 1, mutualist) the susceptibility of the host to patch-level extinction events. When *σ* = 1, the microbe has no effect on host survival at the patch scale. Notice that equation (3) can be reduced to a system of two equations using the relationship 1 = *E* + *P*_0_ + *P*_*m*_.

### Host-scale Metapopulation Model with Rescue Effect

In the previous model, we did not consider variation in host abundance across host habitat patches. By making this approximation, we eliminated the potential for a ‘rescue effect’ – a commonly studied phenomenon in metapopulations whereby large populations are less likely to go extinct as compared to small populations due to stochastic perturbation.^30^ In classic, single-scale metapopulation models, the rescue effect can impact metapopulation dynamics, including the likelihood of metapopulation persistence.^30,31^ A standard approach for incorporating a rescue effect into the basic Levins model is to consider two different classes of colonized patches – patches with small populations and patches with large populations.^19^ Extending this approach to a multiscale metapopulation model, and specifically focusing on a ‘rescue effect’ only at the host-scale, we define the following system of equations:

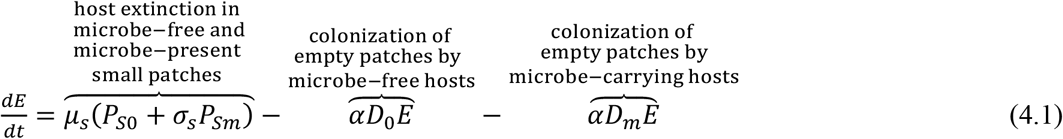

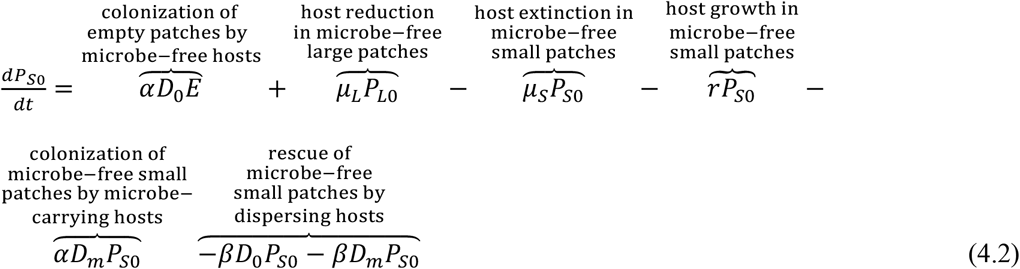

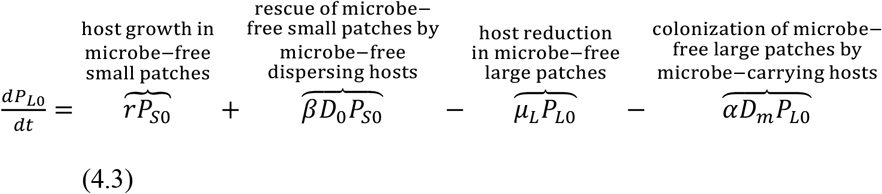

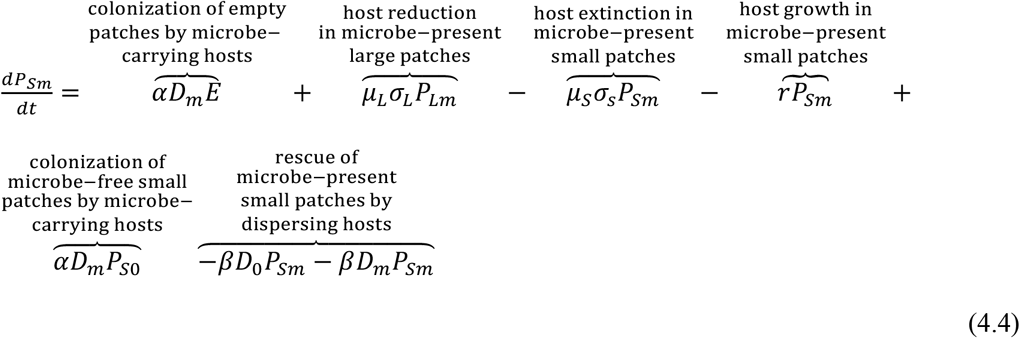

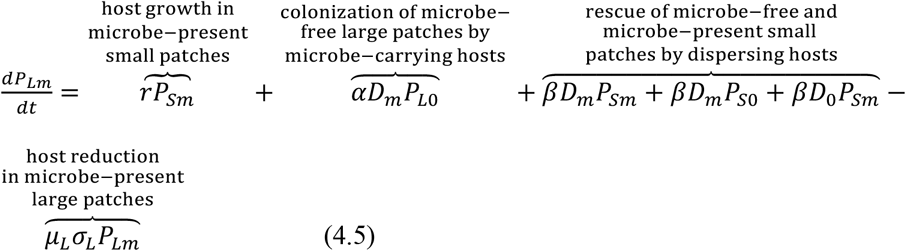

where *E* is the number of empty host patches (no hosts or microbes), *P*_*S*0_ is the number of microbe-free host patches with small host populations (a few hosts present, but none carry the microbe), *P*_*L*0_ is the number of microbe-free host patches with large host populations (many hosts present, but none carry the microbe), *P*_*Sm*_ is the number of microbe-present host patches with small host populations (a few hosts present, and at least some carry the microbe), and *P*_*Lm*_ is the number of microbe-present host patches with large host populations (many hosts present, and at least some carry the microbe). In equation (4), *µ*_*s*_ is the rate at which host extinction occurs in host habitat patches with small host populations, *µ*_*L*_ is the rate at which the host population dips below the threshold size at which extinction becomes more probable, *r* is the rate at which the host population grows to a size above the threshold for likely host extinction and *β* is the rate at which dispersing hosts ‘rescue’ small host populations by boosting their size over the threshold for host extinction. Finally, we again include the parameters *σ*_*s*_ and *σ*_*L*_, to allow for the possibility that the microbe increases (*σ*_*s,L*_ > 1, pathogen) or decreases (*σ*_*s,L*_ < 1, mutualist) the susceptibility of the host to patch extinction and/or population reduction below the threshold for likely extinction. As in equation (3), equation (4) can be reduced to a system of four equations using the relationship 1 = *E* + *P*_*L*0_ + *P*_*S*0_ + *P*_*Lm*_ + *P*_*Sm*_.

### Linking Microbe- and Host-Scale Dynamics in a Basic Multiscale Metapopulation

We now use equations (1), (2), and (3) to define the number of dispersing hosts that are microbe-free and microbe-carrying respectively. This effectively couples our microbe-scale metapopulation and our host-scale metapopulation. For the basic metapopulation model in equation (3), we assume that each non-empty habitat patch has *k* hosts, where *k* is the carrying capacity of an individual habitat patch. Summing over all habitat patches gives the total host carrying capacity across the metapopulation, *K* = ∑ *k*. If we further assume that there is no difference in the tendency of microbe-free versus microbe-carrying hosts to enter the disperser pool (an assumption that could easily be relaxed, but at the expense of additional parameters), then the rate of change of microbe-free and microbe-carrying dispersers is given by:

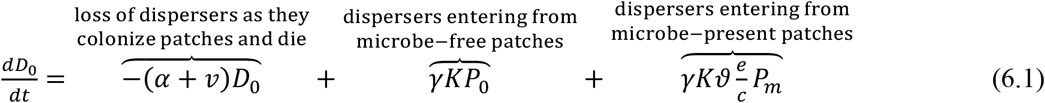

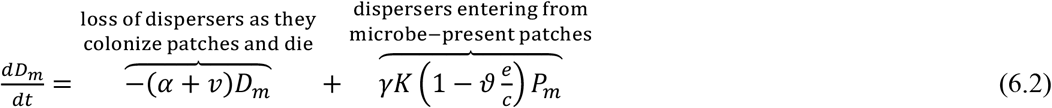

where *v* is the rate of disperser mortality and *γ* is the rate at which hosts located within a patch exit the patch and become dispersers. In equation (6), we have assumed a separation of timescales, such that the microbe, once introduced into a patch, rapidly reaches an equilibrium distribution across hosts (i.e., a pseudo-steady state approximation, see Appendix 1). Under this assumption, equation (2) gives the fraction of microbe-carrying and microbe-free hosts present in a microbe-present patch as 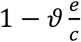 and 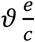 respectively. From equation (6), we find the equilibrium number of microbe-free and microbe-carrying dispersers as:

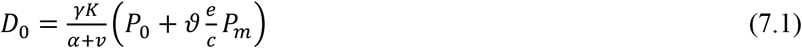

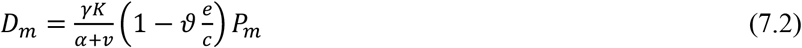

Substituting equation (7) into equation (3) gives (see Supplementary File I):

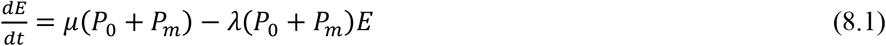

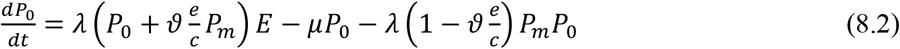

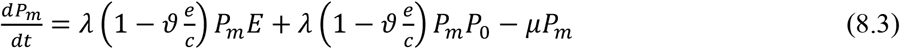

where 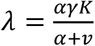 is an effective rate of colonization of empty host habitat patches by hosts from other occupied habitat patches.

### Linking Microbe- and Host-Scale Dynamics in a Metapopulation with a Rescue Effect

For the metapopulation model with a host-scale rescue effect (equation (4)), we take a similar approach. Specifically, we assume that each habitat patch with a large host population (*P*_*Lm*_, *P*_*L*0_) has *k* hosts, where *k* is the carrying capacity of a habitat patch and *K* = ∑ *k* is the total carrying capacity across the entire host metapopulation. For habitat patches with small host populations (*P*_*Sm*_, *P*_*S*0_), we assume that the patch contributes a negligible number of hosts to the disperser pool. In this case, equation (8) can again be used to describe *D*_0_ and *D*_*m*_ by replacing *P*_0_ and *P*_*m*_ with *P*_*L*0_ and *P*_*Lm*_ respectively. Substituting this modified version of equation (8) into equation (4) gives the metapopulation model with a rescue effect (see Supplementary File II).

### Extension to a Mainland-Island Metapopulation

One common extension of single-scale metapopulation models is the ‘mainland-island model’. This model assumes that dispersers come not only from small, isolated habitat patches, but also, from a more permanent mainland population. Importantly, the mainland introduces a constant and stable source of dispersers into the metapopulation. This is because the population on the mainland is assumed to be large enough to avoid population fluctuations and extinction due to stochastic events. To account for a mainland, we can augment equation (7) as follows:

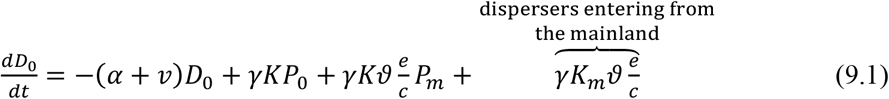

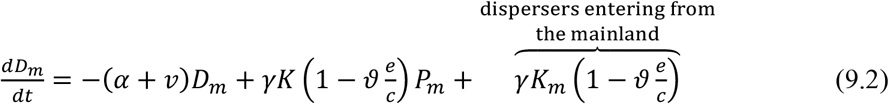

where *K*_*m*_ is the size of the mainland population and *K* is, as before, the size of the population across the smaller ‘islands’ comprising the metapopulation. From equation (9), we find the equilibrium number of microbe-free and microbe-carrying dispersers as:

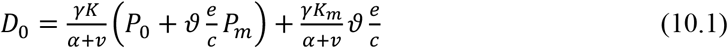

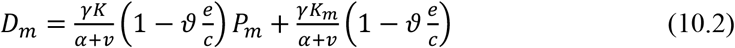

Notice that, when the size of the mainland population is much larger than the size of the island populations, equation (10) reduces to the constants 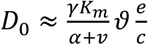 and 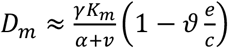 respectively. Substituting equation (10) into equation (3) gives the mainland-island version of the basic multiscale metapopulation model (see Supplementary File III).

In what follows, we analyze our three models (equations (3), (4) and (9)), both by solving for equilibria and determining stability criteria and through numerical simulation. For most analyses, we use Maple 2024 to solve for solutions and perform numerical simulations. The sole exception is for the bifurcation diagrams, where we use the MatCont 7.1 Continuation Toolbox in MATLAB to perform numerical continuation on our sets of ordinary differential equations.

## Results

### Basic Multiscale Metapopulation Model with a Neutral Microbe

We begin by considering the basic metapopulation model (equation (8)). Assuming that the microbe is neutral in its effect on both microbe-scale (*ϑ* = 1) and host-scale (*σ* = 1) extinction, there are four possible equilibrium solutions to equation (8). Of these, three are biologically realistic (i.e., non-negative for at least some parameters). We designate these three solutions as follows based on their biological interpretations:

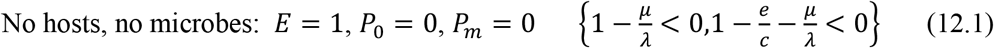

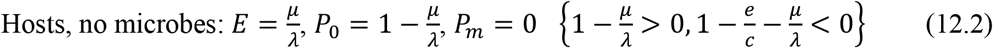

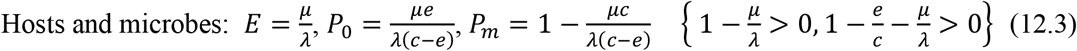

where the linear stability criteria for each of the three solutions are shown in braces (see Supplementary File I). Notice that, for any finite *e/c* > 0, the solution with microbes present (equation 12.3) will always become unstable before the solution with only hosts (equation 12.2); thus, the microbe always go extinct before the host. Further, notice that the persistence criterion for hosts (equation 12.2) remains unchanged from a basic Levins metapopulation model at the host-scale. That is, the microbe does not impact host persistence. The host does, however, impact microbial persistence. In particular, whereas the microbial persistence criterion in a single microbe-scale metapopulation is 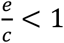, the microbial persistence criterion in our multiscale metapopulation model is 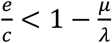. Thus, disturbance at the host patch scale reduces microbial persistence. Said differently, in a multiscale metapopulation, microbial colonization of new hosts must not only exceed microbial loss from already colonized hosts but must do so by a sufficient margin to balance additional losses that stem from host patch extinction events.

In Figure 2, we show predictions for the fraction of patches that are occupied by hosts with and without microbes for a variety of different combinations of colonization:extinction rates at the microbe (*c/e*)- and host (*λ/µ*)-scales. In Figure 2A, for example, we show a scenario with *c/e* < 1, meaning that, even based on a single microbe-scale Levins model, the microbe should not persist. As expected, in this scenario the host occupies patches in accordance with a host-scale Levins model, while the microbe is absent from the system. By contrast, in Figure 2B, we show a scenario with *c/e* > 1. Here, the microbe should persist based on a single microbe-scale Levins model. However, for the range of host colonization:extinction rates considered, we again see that the host occupies patches according to a host-scale Levins model while the microbe is absent. In this case, the added disturbance caused by host-scale extinctions prevents the microbe from establishing even when it could otherwise. In Figures 2C and 2D, we show scenarios with *c/e* ≫ 1. In both cases, the microbe can persist, although only when host colonization:extinction rates are substantially higher than the threshold *λ/µ* > 1 required for host persistence.

**Figure 2.**
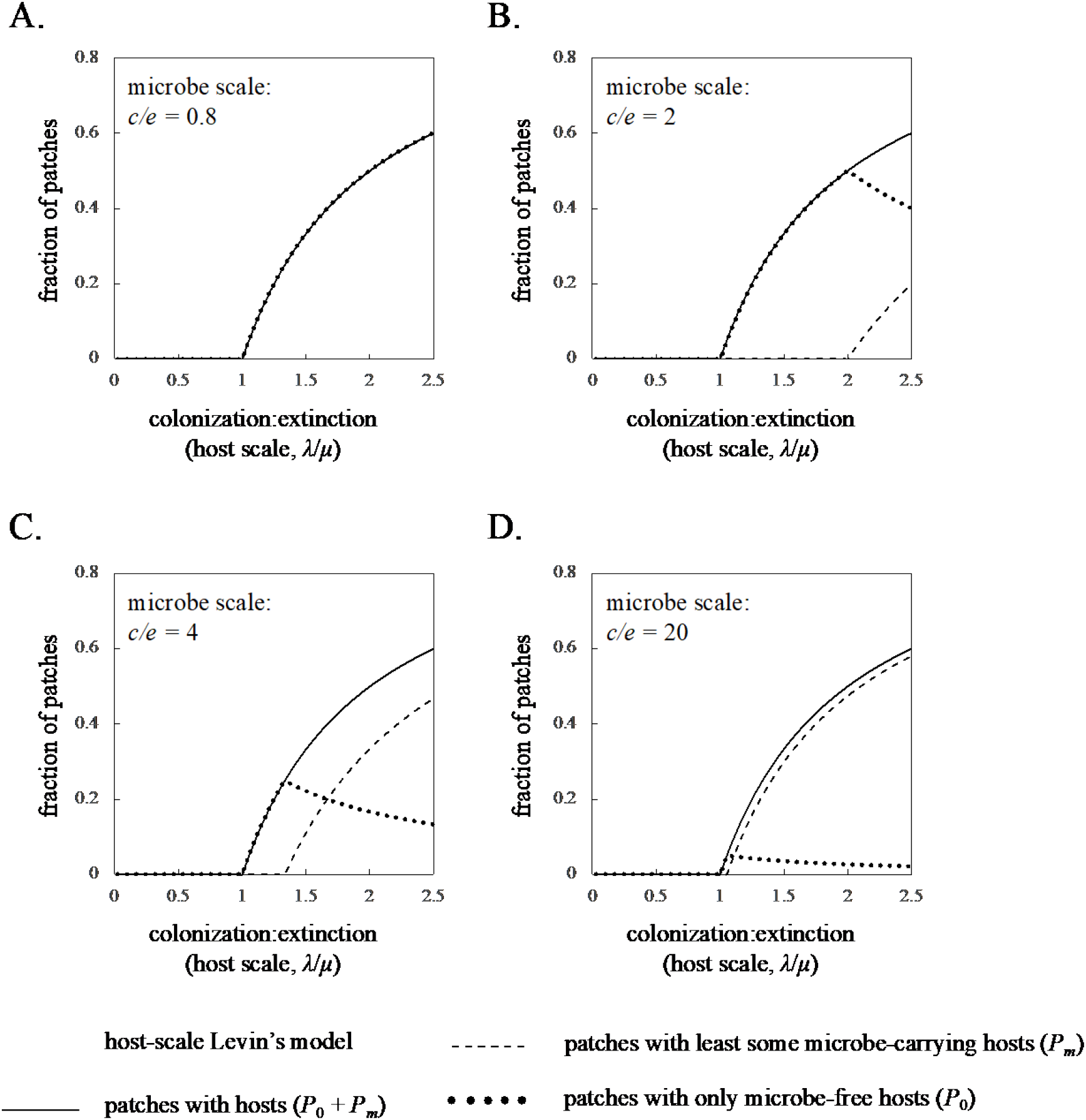
Basic multiscale metapopulation model predictions for a neutral (σ = 1) microbe. Panels show the fraction of patches occupied by all hosts (solid line), only hosts without the microbe (dotted line) and at least some hosts that carry the microbe (dashed line) as a function of host colonization:extinction rates for microbe colonization:extinction rates of (A) 0.8, (B) 2, (C) 4 and (D) 20. In all panels, the thick grey line is the prediction for patch occupancy of the host based on the host-scale Levins model.

### Basic Multiscale Metapopulation Model with a Pathogenic and/or Beneficial Microbe

Setting *σ* ≠ 1 and/or *ϑ* ≠ 1 in equation (8) results in somewhat more complicated expressions for the fraction of empty and occupied patches but only when the microbe persists. Specifically, the equilibrium solutions to equation (8) become:

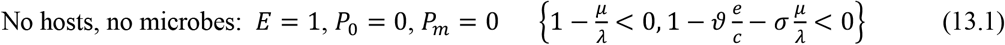

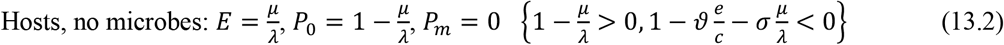

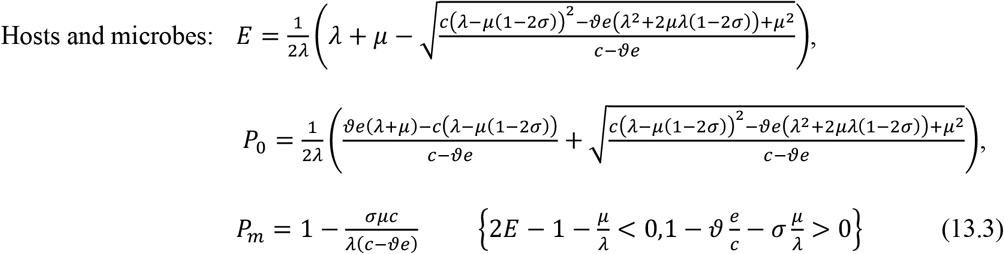

where again, we show the linear stability criteria for each solution in braces (see Supplementary File I). As expected, *σ* and *ϑ* only affect patch occupancy when the microbe persists (equation (13.3)). Both *σ* and *ϑ* can also, however, alter the persistence criteria themselves. For microbial persistence, the new criterion becomes 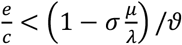. Thus, as compared to a neutral microbe, a pathogenic microbe (*σ* > 1, *ϑ* > 1) must colonize hosts faster or be lost from hosts more slowly in order to persist. Meanwhile, a beneficial microbe (*σ* < 1, *ϑ* < 1) can persist even at slower host colonization rates and faster host clearance/turnover rates. For host persistence, the effects of *σ* and *ϑ* are slightly more complicated. For a pathogenic microbe (*σ* > 1, *ϑ* > 1) and for any finite *e/c* > 0 and/or finite 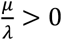, the solution with microbes present (equation (13.3)) will always become unstable before the solution with only hosts (equation (13.2)). Consequently, the pathogen cannot alter the persistence criterion for the host. However, for a beneficial microbe (*σ* < 1, *ϑ* < 1), this is not the case. Rather, for a beneficial microbe, it is possible that 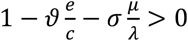 even when 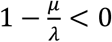. More specifically, this will occur when 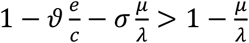 which simplifies to 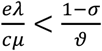. In this case, the beneficial microbe enables the host to persist at lower host patch colonization:extinction ratios than would otherwise be possible (compare Figure 4A to Figures 4C and 4D). Further, the stable solution goes directly from being the host and microbe solution (equation (13.3)) to the no host, no microbe solution (equation (13.1)). Thus, the microbe persists until host extinction, at which point both the host and the microbe are lost simultaneously.

Figures 3 and 4 show predictions for the fraction of patches that are occupied by hosts with and without microbes for a variety of different combinations of colonization:extinction rates at the microbe (*c/e*)- and host (*λ/µ*)-scales for pathogenic (*σ* > 1, *ϑ* = 1; Figure 3) and beneficial (*σ* < 1, *ϑ* = 1; Figure 4) microbes respectively. As in Figure 2, when *c/e* < 1, the microbe goes extinct, and the host occupies patches according to a classic Levins model. Thus, as with a neutral microbe, the minimum criterion for the microbe’s persistence in a multiscale metapopulation is the microbe’s persistence in a single, microbe-scale metapopulation. When *c/e* > 1, beneficial and pathogenic microbes have the potential to persist. However, as with neutral microbes, this depends on whether the microbe can tolerate the added disturbance due to host patch destruction. Notably, a beneficial microbe can persist at lower microbe colonization:extinction rates and lower host colonization:extinction rates than either a neutral microbe or a pathogen (compare, for example, Figure 4B to Figures 2B and 3B). Meanwhile, a pathogenic microbe cannot persist even at host colonization:extinction rates that enabled persistence of a neutral microbe (compare, for example, Figs. 3C and 3D to Figs. 2C and 2D).

**Figure 3.**
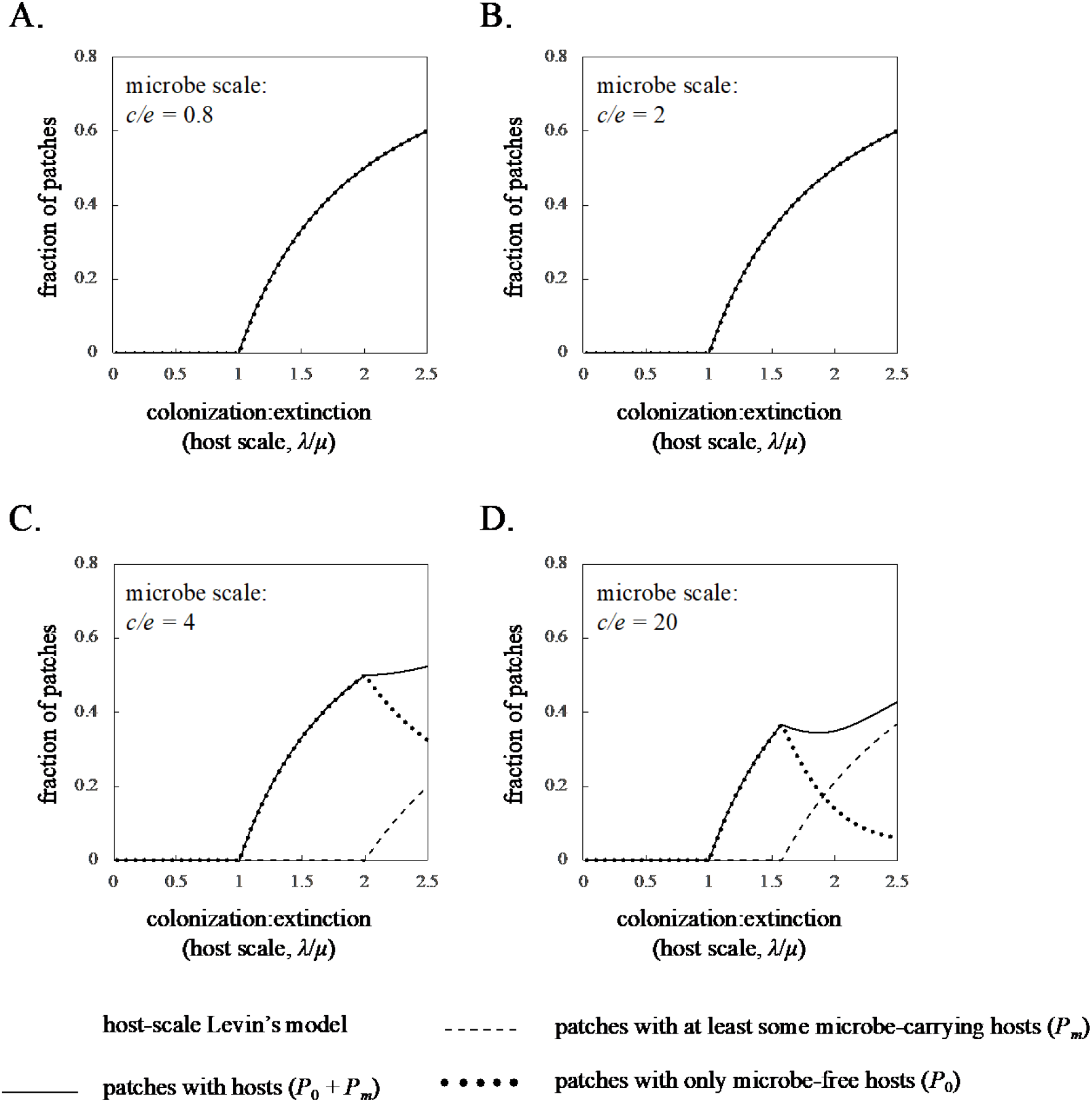
Basic multiscale metapopulation model predictions for a pathogenic (σ = 0.5) microbe. Panels show the fraction of patches occupied by all hosts (solid line), only hosts without the microbe (dotted line) and at least some hosts that carry the microbe (dashed line) as a function of host colonization:extinction rates for microbe colonization:extinction rates of (A) 0.8, (B) 2, (C) 4 and (D) 20. In all panels, the thick grey line is the prediction for patch occupancy of the host based on the host-scale Levins model.

**Figure 4.**
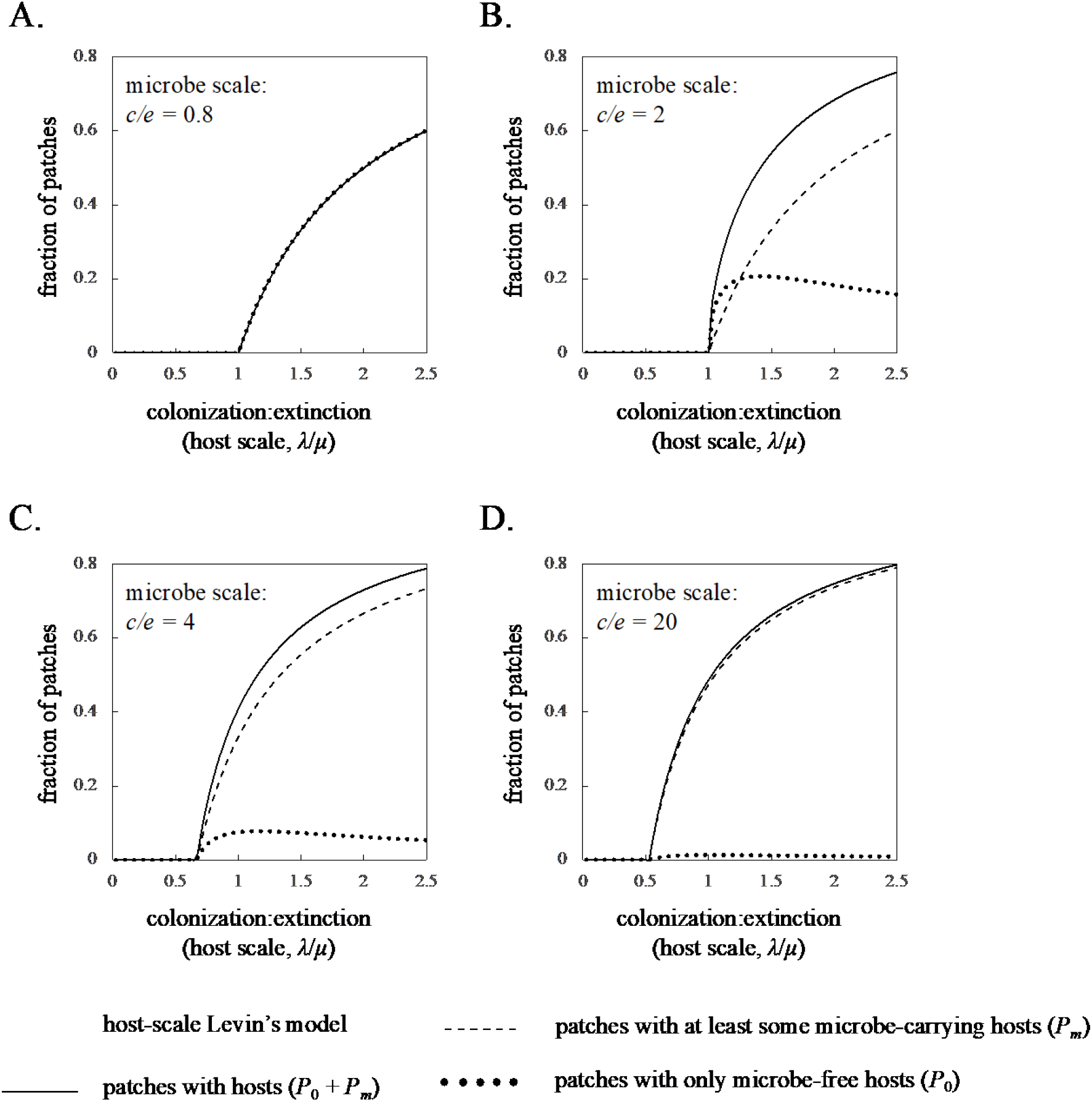
Basic multiscale metapopulation model predictions for a beneficial (σ = 1.5) microbe. Panels show the fraction of patches occupied by all hosts (solid line), only hosts without the microbe (dotted line) and at least some hosts that carry the microbe (dashed line) as a function of host colonization:extinction rates for microbe colonization:extinction rates of (A) 0.8, (B) 2, (C) 4 and (D) 20. In all panels, the thick grey line is the prediction for patch occupancy of the host based on the host-scale Levins model.

Unlike neutral microbes, when pathogenic or beneficial microbes do persist, they impact host occupancy in the metapopulation. Thus, for example, when a pathogen persists (Figures 3C and 3D), there is an overall reduction in the number of occupied host patches as compared to what would be expected based on a simple host-scale Levins model or a multiscale metapopulation model with a neutral microbe. Likewise, when a beneficial microbe persists, there is an overall increase in the number of occupied host patches as compared to a simple host-scale Levins model or a multiscale metapopulation model with a neutral microbe. Finally, as suggested by our analysis of stability criteria, the pathogen cannot impact host persistence, because the microbe is always driven extinct before the host (see Figure 3). By contrast, the beneficial microbe can and does impact host persistence, allowing the host to persist at lower host colonization:extinction ratios than predicted based on the simple, host-scale Levins model (see Figure 4).

### Multiscale Metapopulation Model with a Host Rescue Effect

Closed form, equilibrium solutions for the multiscale metapopulation model with a host rescue effect are not possible. Thus, we resort to numerical solutions (see Supplementary File II). Figure 5 shows predictions for the equilibrium fraction of patches that are occupied by hosts with and without microbes for a variety of different combinations of colonization:extinction ratios at the microbe (*c/e*)- and host (*λ/µ*)-scales for neutral (Figure 5A-D), pathogenic (Figure 5E-H) and mutualistic (Figure 5I-L) microbes respectively. Comparing Figure 5 to Figures 2-4, we see that a rescue effect enables both higher host and higher microbe occupancy rates for any given colonization:extinction ratio at the microbe (*c/e*)- or host (*λ/µ*)-scale. Likewise, a rescue effect enables persistence of both the microbe and the host at lower microbe (*c/e*)- and host (*λ/µ*)-colonization:extinction ratios. However, as in classic, single-scale metapopulation models with a rescue effect^19^, this comes at the cost of bistability. Thus, there emerges a region, defined by a range of microbe (*c/e*)- and host (*λ/µ*)-colonization:extinction ratios, where two different host or host and microbe occupancies levels are possible, depending on initial conditions.

**Figure 5.**
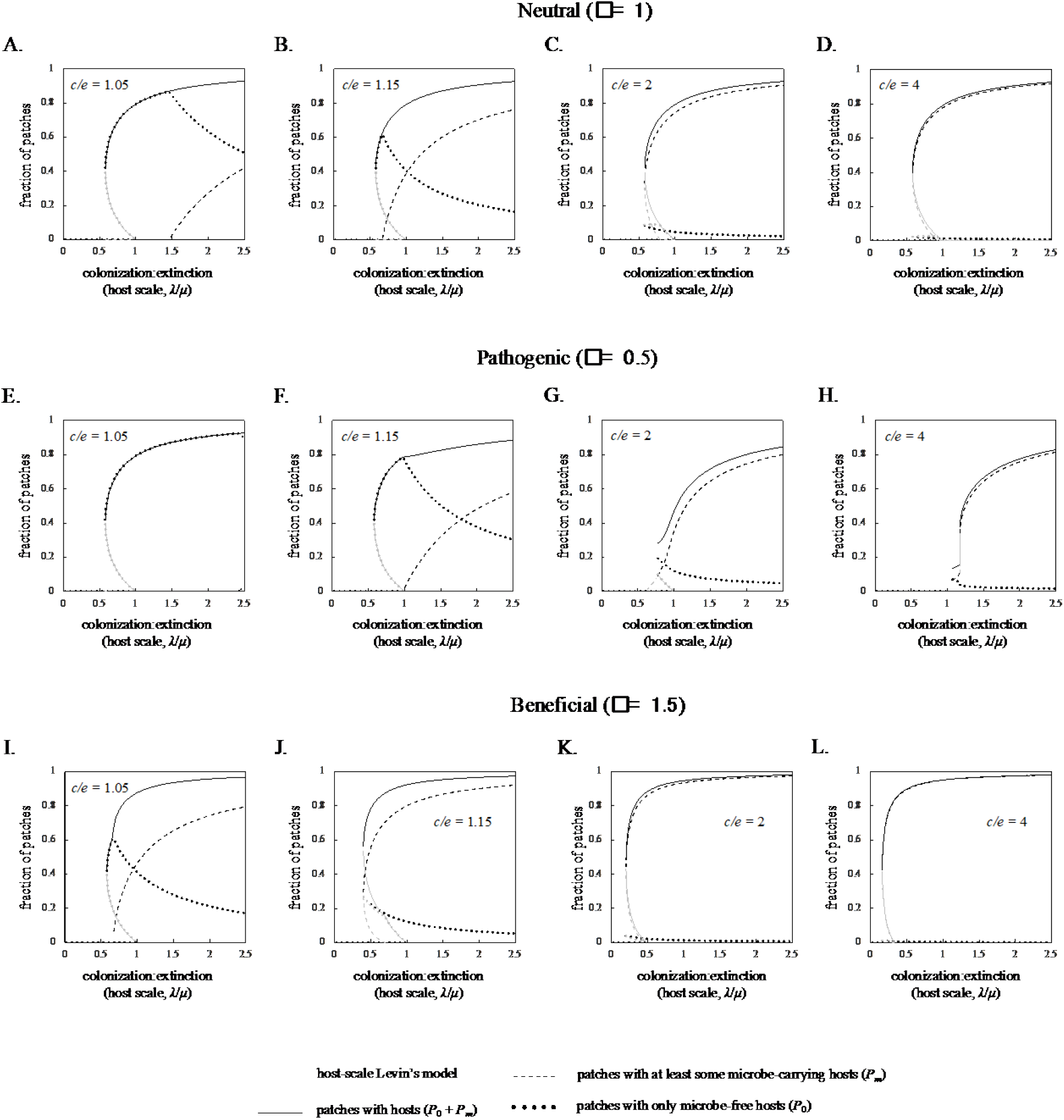
Multiscale metapopulation model predictions for a neutral (σ = 1, A-D), pathogenic (σ = 0.5, E-H) and mutualistic (σ = 1.5, I-L) microbe with a host rescue effect. Panels show the fraction of patches occupied by all hosts (solid line), only hosts without the microbe (dotted line) and at least some hosts that carry the microbe (dashed line) as a function of host colonization:extinction rates for microbe colonization:extinction rates of (A,E,I) 1.05, (B,F,J) 1.15, (C, G, K) 2 and (D,H,L) 4. In all panels, thin black lines are used to indicate stable fixed-point solutions, while thin grey lines are used to indicate unstable fixed point solutions. The thick grey line is the prediction for patch occupancy of the host based on the host-scale Levins model.

Interestingly, in our multiscale metapopulation model, the persistence of the microbe may first occur (i) following the region of host bistability (e.g., Figure 5A), (ii) at some point within the region of host bistability (e.g., Figs. 5B, 5I), or (iii) concomitant with the onset of the region of host bistability (e.g., Figs. 5C,5D, 5J-L), depending on the colonization and extinction dynamics of the microbe itself (*c/e*). Importantly, this means that, for a microbe with a relatively high *c/e* ratio, a rescue effect at the host-scale can enable the microbe to persist until host extinction, even when the microbe is not beneficial to the host. For pathogens, this enables the microbe to destabilize host persistence – a finding that is distinct from the predictions of our basic multiscale metapopulation model without a rescue effect. In that model, the pathogen could reduce host occupancy but could not actually alter host persistence. We see the negative impacts of a pathogen on host persistence in Figures 5E-H. For microbe colonization:extinction ratios of *c/e* = 1.05 and *c/e* = 1.15, the host persists down to a host colonization:extinction ratio of *λ/µ* = 0.581, which is identical to the threshold for host persistence in a simple, host-scale Levins model with a rescue effect. However, at higher microbe colonization:extinction ratios, host persistence is compromised, disappearing at *λ/µ* = 0.625 for *c/e* = 2 and at *λ/µ* = 0.750 for *c/e* = 4. In other words, the pathogen can now make it harder for the host to persist.

Beyond altering host occupancy and persistence criteria, host-associated microbes in systems with host-level rescue effects can have interesting and non-intuitive impacts on system dynamics. In particular, systems with pathogenic microbes and sufficiently large microbe colonization:extinction rates undergo a Hopf bifurcation^32-34^ at low host colonization:extinction ratios. This results in a limit cycle that then disappears at even lower host colonization:extinction rates by way of a homoclinic bifurcation.^34^ Examples of this behavior are shown in Figure 6A and 6B. Notably, the Hopf bifurcation can occur either after the region of bistability (Figure 6A) or within the region of bistability (Figure 6B). Figures 6C and 6D expand on the examples in Figures 6A and 6B by delineating an even wider and more complex range of qualitative dynamics that are possible from the multiscale metapopulation model with a rescue effect. In Figure 6C, for example, we consider the types of dynamics that arise when colonization:extinction ratios at both the microbe (*c/e*)- and host (*λ/µ*)-scales are varied for a pathogenic microbe. Meanwhile, in Figure 6D, we consider the types of dynamics that arise when host colonization:extinction ratios and the impact (from beneficial to pathogenic) of the microbe on the host are varied for a microbe with a relatively high colonization:extinction ratio.

**Figure 6.**
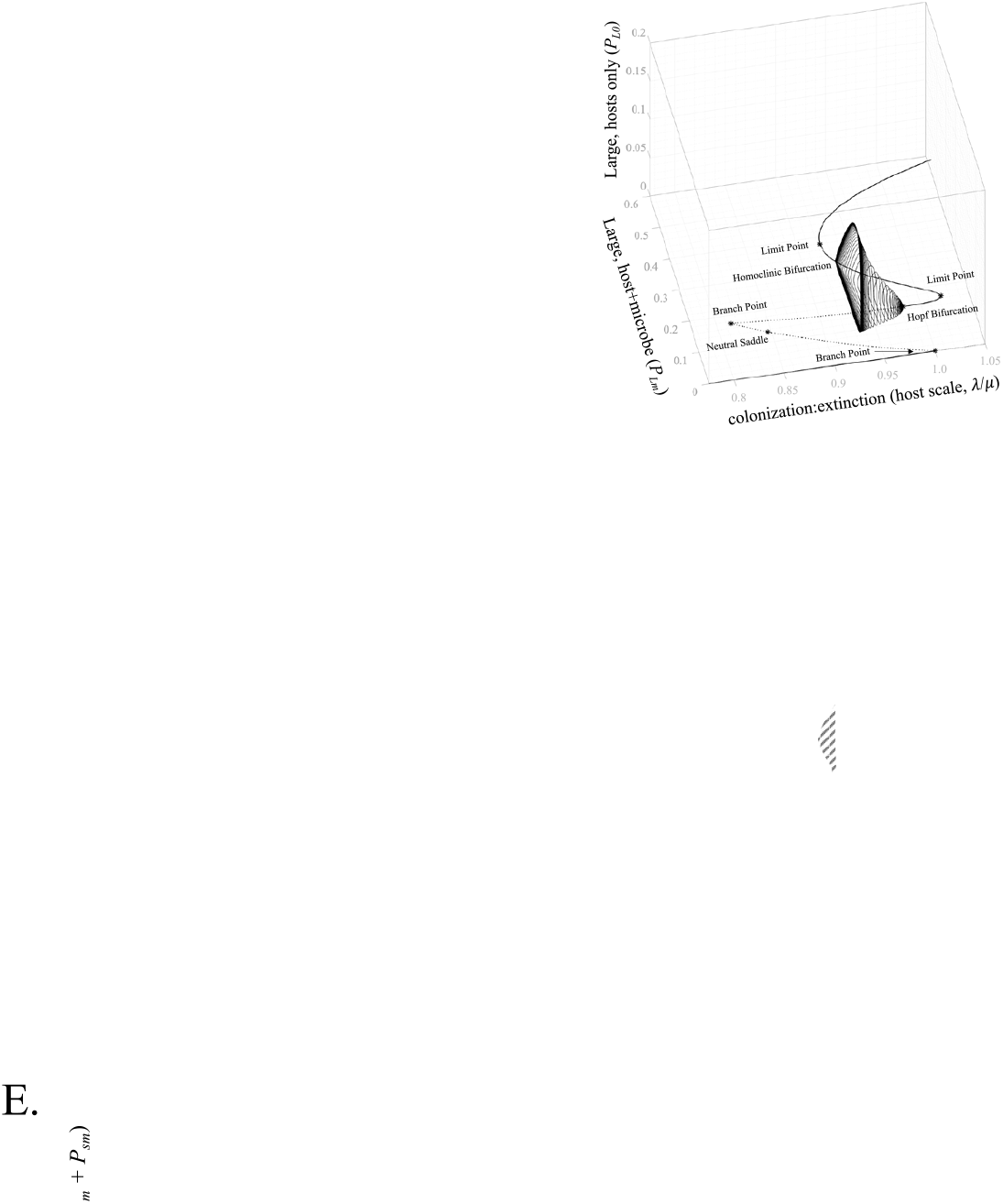
(A,B) Bifurcations and limit cycle dynamics as a function of host colonization:extinction rates for two different pathogenic microbes with differing degrees of pathogenicity. (C,D) Phase diagrams depicting the different types of system dynamics that emerge as a function of host colonization:extinction rates, microbe colonization:extinction rates and microbe pathogenicity/mutualistic effects. Numbered regions exhibit dynamics as follows: (I) host and microbe extinction, (II) bistable with either host and microbe extinction or a host only equilibrium, (III) a host only equilibrium, (IV) bistable with either host and microbe extinction or a host and microbe equilibrium, (V) a host and microbe equilibrium, (VI) either host and microbe extinction or a host and microbe limit cycle, (VII) host and microbe limit cycle, (VIII) bistable with either a low occupancy host and microbe equilibrium or a high occupancy host and microbe equilibrium, (IX) either host and microbe extinction or a host and microbe limit cycle or a host and microbe equilibrium, (X) either host and microbe extinction, or a low occupancy host and microbe equilibrium or a high occupancy host and microbe equilibrium. (E) An example of dynamics from the tristable region (IX, *λ* = 0.1905, *c* = 4, σ = 1.3) showing host and microbe extinction (solid grey), a host and microbe limit cycle (dashed black) or a host and microbe equilibrium (solid black), depending on initial conditions.

### Mainland-Island Multiscale Metapopulation Model

For the extension to the basic mainland-island multiscale metapopulation model, there is a single equilibrium solution with both the microbe and the host present (see Supplemental Information III):

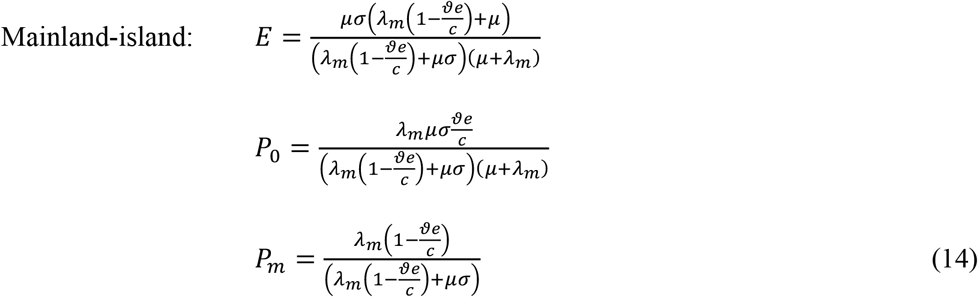

From equation (14), we see that microbe persists whenever 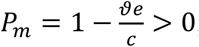, while the host persists whenever *λ*_*m*_ > 0. Thus, as expected, hosts persist in the metapopulation for any finite host immigration rate from the mainland, while microbes persist in the metapopulation provided that they can also persist on the mainland. Figure 7 reiterates these findings, showing predictions for the fraction of patches that are occupied by hosts with and without microbes for a variety of different combinations of colonization:extinction rates at the microbe (*c/e*)- and host (*λ/µ*)-scales.

**Figure 7.**
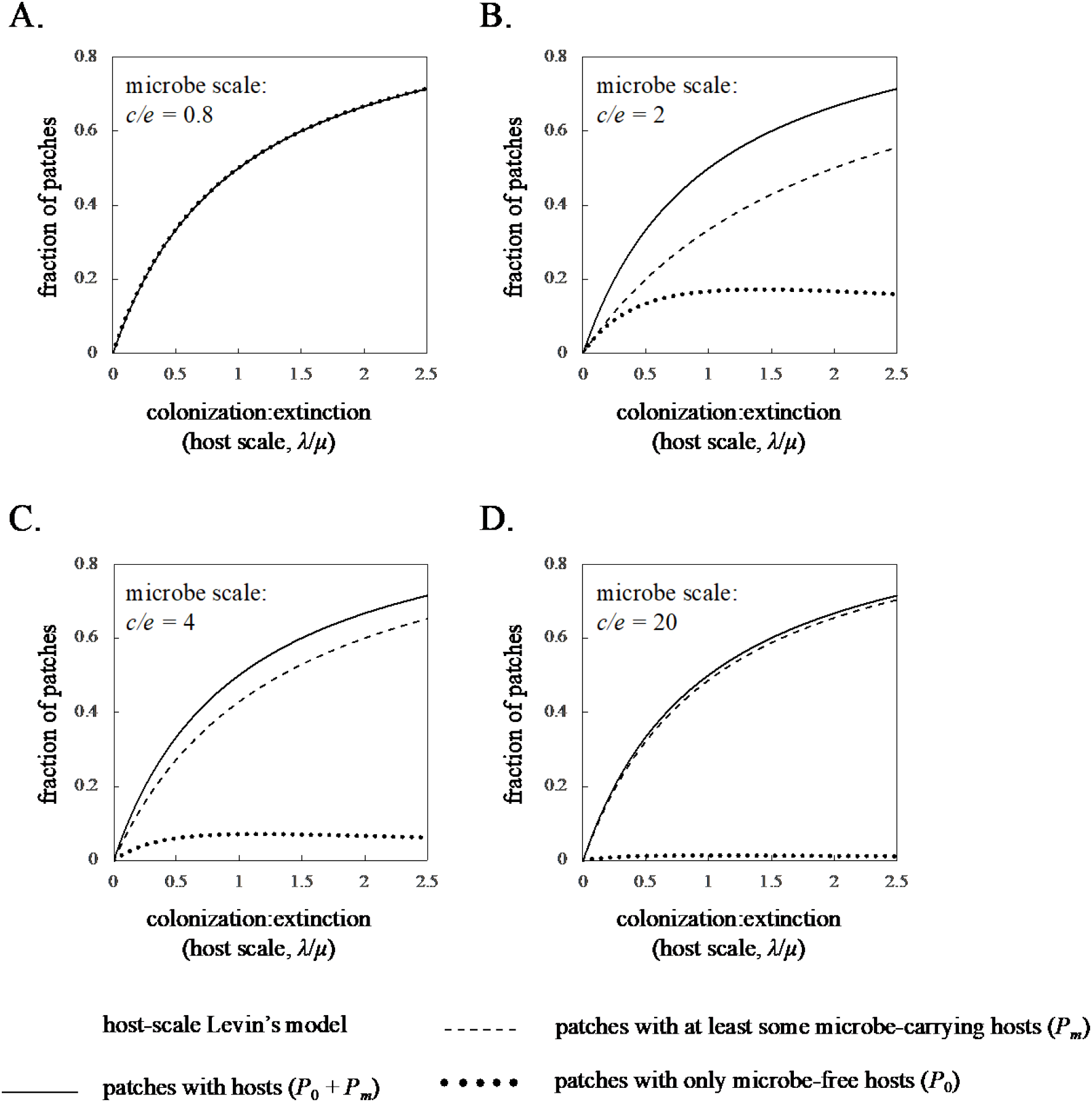
Mainland-island multiscale metapopulation model predictions for a neutral (σ = 1) microbe. Panels show the fraction of patches occupied by all hosts (solid line), only hosts without the microbe (dotted line) and at least some hosts that carry the microbe (dashed line) as a function of host colonization:extinction rates for microbe colonization:extinction rates of (A) 0.8, (B) 2, (C) 4 and (D) 20. In all panels, the thick grey line is the prediction for patch occupancy of the host based on the host-scale mainland-island Levins model.

## Discussion

In this paper, we use an extension to the classic Levins model to explore properties of multiscale metapopulations and to contrast multiscale metapopulations to comparable metapopulations at a single scale. Importantly, our study includes full colonization and extinction dynamics at two scales. By contrast, similar studies on patchy SIS models include colonization at by the host and microbe at the host and microbe scales respectively, as well as extinction of the microbe at the microbe-scale. In general, however, they do not include extinction of the host at the host-scale (but see Hess 1996 and 1999^11,12^ for several studies where patch destruction was included in an SI model for disease). Whereas patchy SIS models are good descriptions for disease spread in a system with dispersal limitation of the host, our model captures both dispersal limitation of the host and host patch destruction within an SIS framework. This is important for systems where host patch destruction plays a role in determining the prevalence and persistence of host-associated organisms, for example ranavirus or chytrid infection in green salamanders^35-37^ or disease spread in butterfly species inhabiting ephemeral environments.^38,39^ Beyond inclusion of host patch destruction, our model is also more general than typical patchy SIS models in that it considers not only pathogens but also beneficial host-associated organisms. Overall, we find frequent deviations between multiscale and single scale model predictions. This includes deviations in persistence thresholds, equilibrium patch occupancy and qualitative system dynamics. However, microbe and host models are impacted differently by the extension to two scales. In what follows, we discuss how multiscale dynamics impact host and microbe model predictions, focusing particularly on the interplay between host colonization:extinction dynamics and microbe colonization:extinction dynamics.

At the microbe-scale, host-associated organisms experience both their own metapopulation structure, as well as the metapopulation structure of the host. Thus, to persist, host-associated organisms must combat both extinction due to loss of individual hosts, and extinction due to loss of entire host populations. From the microbes’ perspective, the distinction between these two forms of host loss is largely irrelevant. Both individual and population-level host loss result in extinction of microbial patches. Consequently, the two colonization:extinction ratios - *e/c* at the microbe-scale and *µ/λ* at the host-scale – enter the microbe persistence criterion additively. The additive nature of the two colonization:extinction ratios on microbial persistence has several consequences. First, it means that a minimum criterion for persistence of the microbe in the host metapopulation is persistence of the microbe in a single host population (see Equations (12) and (13)). In other words, host patchiness cannot cause the microbe to persist when it would not persist in a single, well-mixed host population – a finding consistent with classic patchy SIS models^40^ (but see Wang and Mulone 2004 and Allen et al. 2007^21,41^ for exceptions). Second, because the two colonization:extinction ratios appear additively in the microbe persistence criterion, the microbe cannot persist without persistence of the host. Though this should be obvious for any obligately host-associated organism, recognizing the dependence of microbe persistence on host persistence is helpful for interpreting the effects of pathogens on host metapopulations (see below). Finally, the additive nature of the two colonization:extinction ratios on microbe persistence means that full host metapopulation structure – that is, including both host dispersal limitation *and* host patch extinction - not only fails to promote microbial persistence but actually inhibits it. In particular, any level of disturbance to host patches (i.e., *µ* > 0) uniformly lowers microbe persistence in the host metapopulation relative to microbe persistence in a single, undisturbed host population.

Whereas host-scale metapopulation dynamics always have a negative impact on microbe persistence, the effects of microbe-scale metapopulation dynamics on the host are more subtle and context dependent. By definition, a neutral microbe can have no effect on the host. Thus, it should come as no surprise that neutral microbes neither increase nor decrease host persistence or host patch occupancy (see Figure 2, Figure 5A-D). Rather, the neutral microbe is a passenger whose persistence and occupancy is affected by host metapopulation structure but whose own metapopulation structure has no effect on the host. This is not the case, however, for either pathogens or beneficial microbes. Across all models, pathogens lower host patch occupancy (see Figure 3, Figure 5E-H). In some cases, this leads to the counterintuitive result that lower host colonization:extinction ratios lead to higher host patch occupancy (see, for example, Figure 3D). This occurs because the microbe experiences both scales of metapopulation structure and is thus always more limited than the host. As a consequence, the pathogen suffers disproportionately from high host disturbance regimes. Coupled to the nonlinear effects of the colonization:extinction ratios on patch occupancy in Levins models, the net result is a range of host colonization:extinction ratios where pathogen occupancy falls off much faster than host occupancy, which then benefits the host. While pathogens generally lower host patch occupancy, they can only do so in host metapopulations where they persist. Notably, in the simplest model without a rescue effect, pathogens cannot persist down to the point of host extinction. This is because the host only faces colonization:extinction dynamics at the host-scale, whereas the pathogen faces the same colonization:extinction dynamics at the host-scale and then additional colonization:extinction dynamics at the microbe-scale. Consequently, persistence of the pathogen is always more limited than persistence of the host. Importantly, because the pathogen cannot persist at the host extinction threshold, the pathogen cannot impact host persistence. Instead, the pathogen will always go extinct before the host and, once the pathogen goes extinct, the host follow a classic single-scale Levins metapopulation at the host-scale.

Like pathogens, beneficial microbes also alter host patch occupancy. However, in the case of beneficial host-associated organisms, they increase, rather than decrease patch occupancy. Further, unlike pathogens, beneficial microbes can alter host persistence, even in the simplest case of a system without a host rescue effect. More specifically, beneficial microbes can lower the host persistence threshold if the beneficial microbe either significantly reduces individual host death (or microbial loss from individual hosts, *ϑ* ≪ 1) or if the beneficial microbe significantly reduces host patch destruction (*σ* ≪ 1). A beneficial microbe is also more likely to alter host extinction thresholds when the system is characterized by high microbe-scale colonization:extinction ratios and/or low host-scale colonization:extinction ratios (see Equation (13)). Though somewhat less intuitive, these conditions emerge because the microbe persistence criterion is equal to the host persistence criterion plus an additive constraint based on individual host death (or microbial clearance from an individual host). As a result, minimizing host death (or microbial clearance) relative to host patch destruction forces the host and microbe persistence criteria closer together. Said differently, a microbe that is primarily facing extinction due to the same dynamics that threaten the host is more likely to persist at the limits of host persistence and it is in this region where a beneficial microbe can have its greatest positive effects on host persistence.

Perhaps one of the most interesting, but also alarming, findings from our model is how a host rescue effect impacts the dynamics of the multiscale system. First introduced by Brown and Kodric-Brown^30^, the idea of a rescue effect is that, in metapopulations with numerous large local populations, there are a wealth of dispersers that migrate to small local populations and, in doing so, ‘rescue’ the small local populations from imminent extinction. By contrast, in metapopulations with only a few small local populations, there are fewer dispersers, and many of the dispersers that do exist are wasted on empty patches where they do not establish or in small local populations that go extinct anyway. As in single-scale metapopulation models, a host rescue effect creates a region of bistability where, for the same colonization:extinction ratios, large metapopulations stay large but small metapopulations go extinct. In our system, this creates a scenario where a sizeable fraction of host patches may be occupied up to the point where bistability disappears and host extinction becomes the only stable state. Unfortunately, high host occupancy just prior to host extinction provides opportunity for pathogenic host-associated organisms to persist up to the threshold for host persistence. Such persistence then enables pathogens to impact the threshold for host persistence. Thus, in systems where a host rescue effect occurs, the microbe-scale metapopulation can destabilize the host-scale metapopulation, altering both host patch occupancy and host patch persistence. Notably, however, to do so the microbe must still overcome both its own colonization:extinction dynamics and the host colonization:extinction dynamics. Thus, host persistence is only altered at higher microbial colonization:extinction ratios. While a host rescue effect also allows neutral and beneficial microbes to persist at higher rates of host disturbance, the outcomes are less dramatic, since the neutral microbe has no effect on the host under any conditions, and the beneficial microbe was able to increase host persistence even in the simple model without a host rescue. Notably, the host rescue effect does not change the minimum requirement that the microbe be able to persist in a single host population in order to persist in a host metapopulation. However, all else being equal, the host rescue effect does enable the microbe to persist at higher rates of host disturbance.

In addition to altering both host and microbe persistence, a host rescue effect can also have interesting effects on system dynamics. In particular, a host rescue effect can result in stable equilibria giving way to oscillations in both the host and the microbe. However, this only occurs when the microbe is a relatively strong pathogen (see Figure 6D, noticing that limit cycles only appear when *σ* > 1). When the microbe is neutral or beneficial, limit cycles are replaced by simple bistability. In general, pathogen limit cycles occur at intermediate host and microbe colonization:extinction ratios. At lower host colonization:extinction ratios, both the host and the microbe go extinct. At higher host colonization:extinction ratios, there is a single host and microbe equilibrium. At lower microbe colonization:extinction ratios, there is bistability and either the host and microbe go extinct, or there is a host or a host and microbe equilibrium. Finally, at higher microbe colonization:extinction ratios, both the host and the microbe go extinct. Thus, limit cycles appear in a region where the host colonizes new patches slowly enough to attenuate pathogen spread but not slowly enough to drive the pathogen extinct. Meanwhile, the pathogen spreads rapidly enough to persist, but not so rapidly that it drives the host (and by extension itself) extinct. Thus, similar to classic outbreaking disease systems^42^, the limit cycles are driven by overcompensatory dynamics, wherein the pathogen rapidly reduces the host when host patch occupancy is high but then suffers a population crash itself when it cannot find sufficient host patches to colonize. This, then allows host patch occupancy to increase again, repeating a cycle of host growth, followed by pathogen growth, followed by host decline, followed by pathogen decline. In our system, however, oscillations are driven by changes in occupancy across a metapopulation rather than changes in abundances of susceptible hosts, with the precise system parameters that lead to overcompensatory dynamics depending on metapopulation structure at both the microbe and host-scales.

While the goal of this paper was to develop a simple, two-scale metapopulation and to examine the role and interplay of colonization and extinction dynamics at both scales, a number of exciting extensions to this basic model are possible. First, we chose a spatially implicit formulation. Considering spatially explicit models, including both idealized spatial structures like lattices or necklaces, as well as spatial structures based on empirical systems could result in novel outcomes not predicted by our current models. Likewise, we have intentionally focused on a system with identical patches. Work on patchy SIS models, however, has shown that differences in patch size or disease dynamics within patches can result in counterintuitive results. It would be interesting to explore these types of systems within the current modeling framework. Similarly, we have assumed that susceptible and infected hosts show equal propensity for dispersing. By contrast, microbe carrying individuals could disperse more (e.g. because they are healthier or because the microbe manipulates host behavior) or less (e.g., because they are physically compromised) than susceptible individuals. This is an extension that is common in patchy SIS models and could be easily incorporated into Equations (10), albeit with additional parameters. Finally, we have considered a single host and a single microbial taxon. With the advent of sequencing technology, however, it is becoming more and more realistic to track entire communities of host-associated organisms, for example gut or skin microbiomes, across host metapopulations. Extending the current framework to consider microbiomes, rather than single microbial populations, or even to consider microbiomes on entire host communities (i.e., multiple host species) could provide interesting insight into the assembly of microbiomes on hosts that exist in classic metapopulation environments, for example, ephemeral ponds, forest gaps, and fields.

With new and improved tools for tracking microbial constituents on macroscopic hosts – for example qPCR targeting specific pathogens, 16S rRNA gene and ITS sequencing of bacterial and fungal communities and shotgun metagenomics, our ability to study multiscale metapopulations and metacommunities is increasing rapidly. Indeed, recent empirical investigations into real-world microbial metacommunities on host metapopulations suggest interesting patterns which are not well understood or described by current theory. Some members of a microbial metacommunity, for example, are more conserved than others and microbiome similarity across space and time appears to be dependent upon host metapopulation status, though in largely unpredictable ways (Couch et al. 2020). To better describe this emerging body of empirical work, new models are required. Here, we use the classic pillar of metapopulation theory - the Levins model - to begin to develop one such multiscale framework. Ultimately, our model and models like it will provide the theory necessary to guide, describe, predict and explain the structure and dynamics of exciting new multiscale systems that span the vastly different spatial and temporal scales according to which different organisms exist but across which different organisms also interact.

## Supporting information

Supplementary File III

## Acknowledgements/Funding Declaration

None

## Appendix 1: Quasi-steady state approximation

To identify conditions where the quasi-steady state approximation is valid, we focus on the microbe-scale Levin’s model, along with the equations for the host-scale disperser pool:

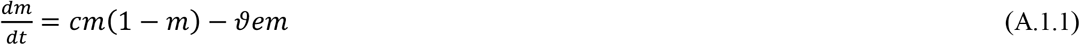

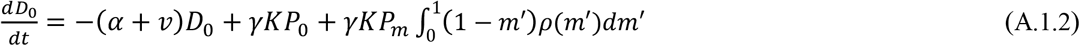

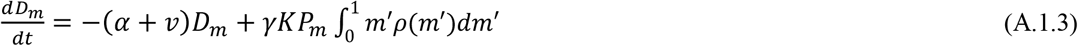

In equation (A.1), we have rewritten the equations for the host-scale disperser pool without making a pseudo-steady state approximation. Specifically, we integrate across all possible proportions of microbe carrying hosts, *m′*, where *ρ*(*m′*) is the fraction of microbe-present patches with *m′* proportion of microbe carrying hosts. We begin by non-dimensionalizing the system in (A.1) using the following relationships:

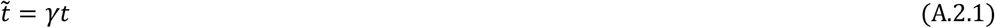

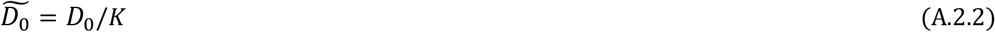

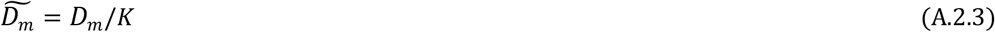

This gives

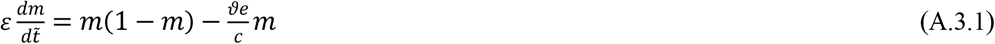

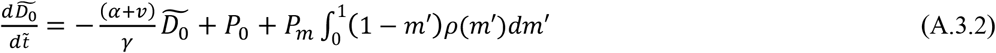

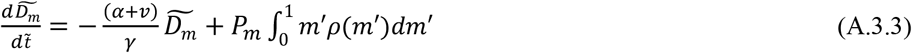

where 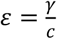. In the event that *γ* ≪ *c*, we can make a pseudo-steady-state approximation on equation (A.3.1). This means that the rate at which hosts leave a patch and enter the disperser pool is significantly slower than the rate at which host-host contact within a patch spreads the microbe from an infected host to a susceptible host. Specifically, taking the limit as *ε* → 0, we find

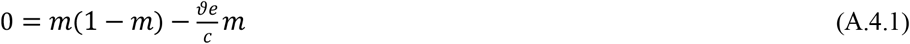

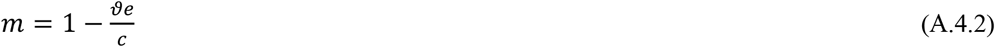

Under this approximation, all microbe-present carrying patches have the same proportion of microbe carrying hosts, thus, 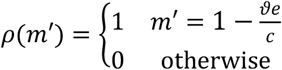. This gives:

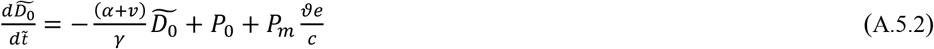

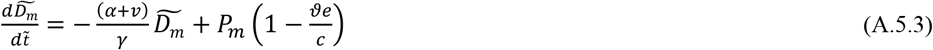

Which are the pseudo-steady state approximation equations given in the main text.

## Appendix 2

Substituting equation (6) into equation (3) and assuming *δ* = 0 and 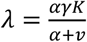gives:

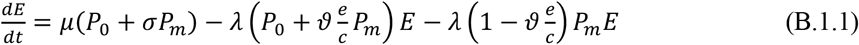

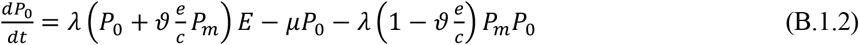

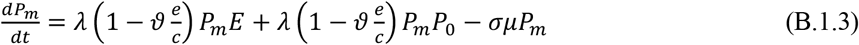

The Jacobian of equation (B.1) is given by

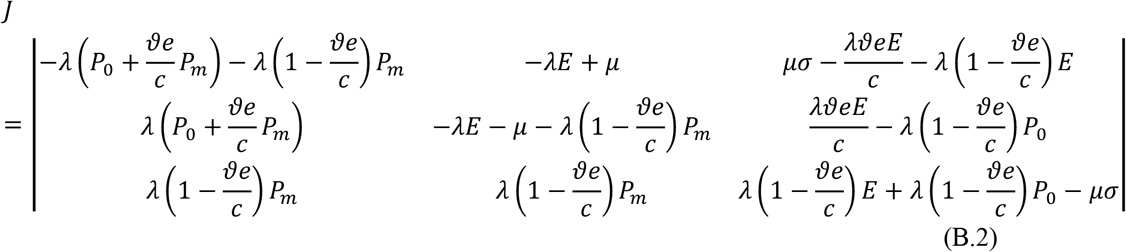

Setting *σ* = 1 and *ϑ* = 1 and solving equation (B.1) gives four possible solutions:

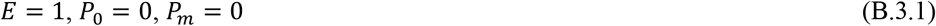

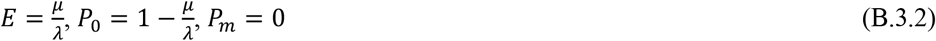

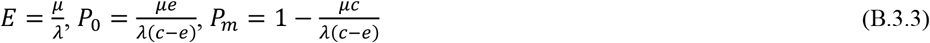

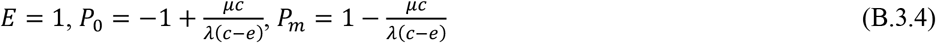

Assuming that all colonization and extinction rates are positive (*µ* > 0, *λ* > 0, *e* > 0, *c* > 0), solution (B.3.1) is always positive, solution (B.3.2) is positive provided *µ* < *λ* (host colonization of empty patches is faster than host extinction from occupied patches) and solution (B.3.3) is positive provided *c* > *e* (microbe colonization of susceptible hosts is faster than microbe extinction from infected hosts) and 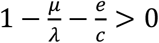. For 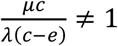, solution (B.3.4) can never be positive since *P*_0_ = −*P*_*m*_.

Substituting equation (B.3.1) into equation (B.2) and solving for the criteria that result in negative eigenvalues gives:

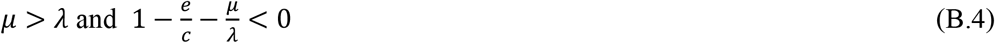

Substituting equation (B.3.2) into equation (B.2) and solving for the criteria that result in negative eigenvalues gives:

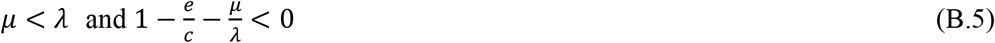

Finally, substituting equation (B.3.3) into equation (B.2) and solving for the criteria that result in negative eigenvalues gives:

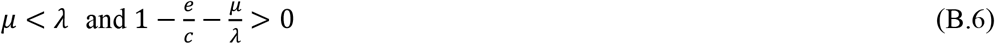

Combining equations (B.4)-(B.6) gives the stability criteria in equation (12) of the main text. An identical approach can be used to solve for the stability criteria in equation (13), although the results are somewhat more complicated, thus we do not show them here (but see Supplementary_File_I.maple) where we use Maple to solve for the eigenvalues of the Jacobian. Likewise, the same approach can be taken to solve for the stability criteria corresponding to the model with the rescue effect (see equation (4)), although in this case it is necessary to solve for stability criteria numerically (see Supplementary_File_II.maple for examples where we do this). Finally, a similar approach can be used to derive stability criteria for the mainland-island model (see Supplementary_File_III.maple).

